# Leveraging family data to design Mendelian Randomization that is provably robust to population stratification

**DOI:** 10.1101/2023.01.05.522936

**Authors:** Nathan LaPierre, Boyang Fu, Steven Turnbull, Eleazar Eskin, Sriram Sankararaman

**Author notes:** These authors contributed equally to this work (Nathan LaPierre, Boyang Fu). Email corresponding authors at and.

## Abstract

Mendelian Randomization (MR) has emerged as a powerful approach to leverage genetic instruments to infer causality between pairs of traits in observational studies. However, the results of such studies are susceptible to biases due to weak instruments as well as the confounding effects of population stratification and horizontal pleiotropy. Here, we show that family data can be leveraged to design MR tests that are provably robust to confounding from population stratification, assortative mating, and dynastic effects. We demonstrate in simulations that our approach, MR-Twin, is robust to confounding from population stratification and is not affected by weak instrument bias, while standard MR methods yield inflated false positive rates. We applied MR-Twin to 121 trait pairs in the UK Biobank dataset and found that MR-Twin identifies likely causal trait pairs and does not identify trait pairs that are unlikely to be causal. Our results suggest that confounding from population stratification can lead to false positives for existing MR methods, while MR-Twin is immune to this type of confounding.

## 1 Introduction

Mendelian Randomization (MR) is a widely-used analytical tool that uses genetic variants (“genetic instruments”) to determine whether one trait (the “exposure”) has a causal effect on another (the “outcome”). With the availability of massive biobank datasets such as the UK Biobank [1], MR analyses have become increasingly powerful and have been used to identify causal relationships between numerous pairs of traits [2, 3, 4, 5, 6]. The validity of MR rests on three key assumptions [7]: (i) that the genetic instrument is significantly associated with the exposure; (ii) that the genetic instrument is independent of confounders of the exposure-outcome relationship; (iii) that the genetic instrument affects the outcome only through its effect on the exposure.

Unfortunately, the latter two assumptions are often violated in practice, due to several factors including horizontal pleiotropy, population stratification (and related phenomena such as assortative mating and dynastic effects), and batch effects. Even when these assumptions are met, the weak effects of typical genetic instruments on the exposure coupled with spurious correlation between genetic instruments and confounders [8] can bias the results of MR analyses (“weak instrument bias”). The problem of population stratification has been extensively studied in the Genome-Wide Association Study (GWAS) literature, and approaches for mitigating its effects have been developed, including the usage of Principal Components Analysis (PCA) and Linear Mixed Models (LMMs) [9]. These approaches have generally been found to be effective at reducing the confounding introduced by population stratification [9]. However, recent studies have demonstrated that, with sample sizes as large as those found in modern biobanks, even a small amount of residual population stratification can cause a considerable amount of bias [10, 11, 12, 13], and may even cause false positives in MR analysis [13, 14]. In addition, while the confounding effects of population stratification are well-known, less attention has been directed towards confounding from other phenomena such as (cros-strait) assortative mating and dynastic effects, which can also cause MR false positives [11, 15]. Recent work has demonstrated that cross-trait assortative mating is widespread and substantially inflates genetic correlation estimates between many trait pairs [16].

It has recently been proposed that family-based genetic datasets could be used in MR studies to avoid confounding from population stratification [11, 17]. A recent suite of methods have been developed for this purpose, and were shown to reduce the bias from this type of confounding [11]. However, like other MR methods, these methods are susceptible to weak instrument bias, which can be substantial for small family-based datasets [11].

In this paper, we introduce MR-Twin, a test for causal effects between pairs of traits that is able to leverage family-based genetic data to provably control for population stratification and utilize publicly-available summary statistics estimated in large biobank datasets to achieve power competitive with top existing methods for the same sample size. MR-Twin tests for a causal effect by comparing an appropriate statistic computed on the offspring in the families to their “digital twins”[18], which are simulated offspring created by sampling offspring genotypes from parental genotypes. By testing for causal effects while conditioning on parental genotypes, MR-Twin avoids confounding caused by population stratification, since offspring genotypes are independent of population information given the genotypes of their parents.

We demonstrate in simulations that MR-Twin robustly controls for confounding resulting from population stratification while other methods yield inflated false positive rates (FPR). We also show that MR-Twin avoids inflated FPR due to weak instrument bias, unlike other family-based methods [11]. Finally, we show that MR-Twin can use trio, parent-child, or sibling genetic data, although only trio data provides complete immunity to confounding from population stratification. In an analysis of 121 trait pairs from the UK Biobank [1], we show that MR-Twin identifies likely causal trait pairs and does not identify trait pairs that are unlikely to be causal. With sizable family-based genetic datasets being increasingly available through resources such as the UK Biobank [1], MR-Twin is a widely-applicable and flexible method that can make the findings of MR analyses more robust. MR-Twin is freely available at https://github.com/nlapier2/MR-Twin.

## 2 Results

### 2.1 Methods Overview

In a Mendelian Randomization (MR) analysis, we wish to determine whether one phenotype (the “exposure”) has a causal effect on another phenotype (the “outcome”) using genetic instrumental variables, which can be either single nucleotide polymorphims (SNPs), a polygenic score, or other genetic features. Under the assumption that the genetic instruments are associated with the exposure and are independent of the outcome given the exposure, the MR effect estimate of the exposure on the outcome will be valid even if there are unobserved confounders of the exposure-outcome relationship. The independence assumption, however, is often violated by population stratification (Figure 1a) or horizontal pleiotropy, as these phenomena cause the genetic instruments to be correlated with the outcome through pathways other than those through the exposure.

**Figure 1:**
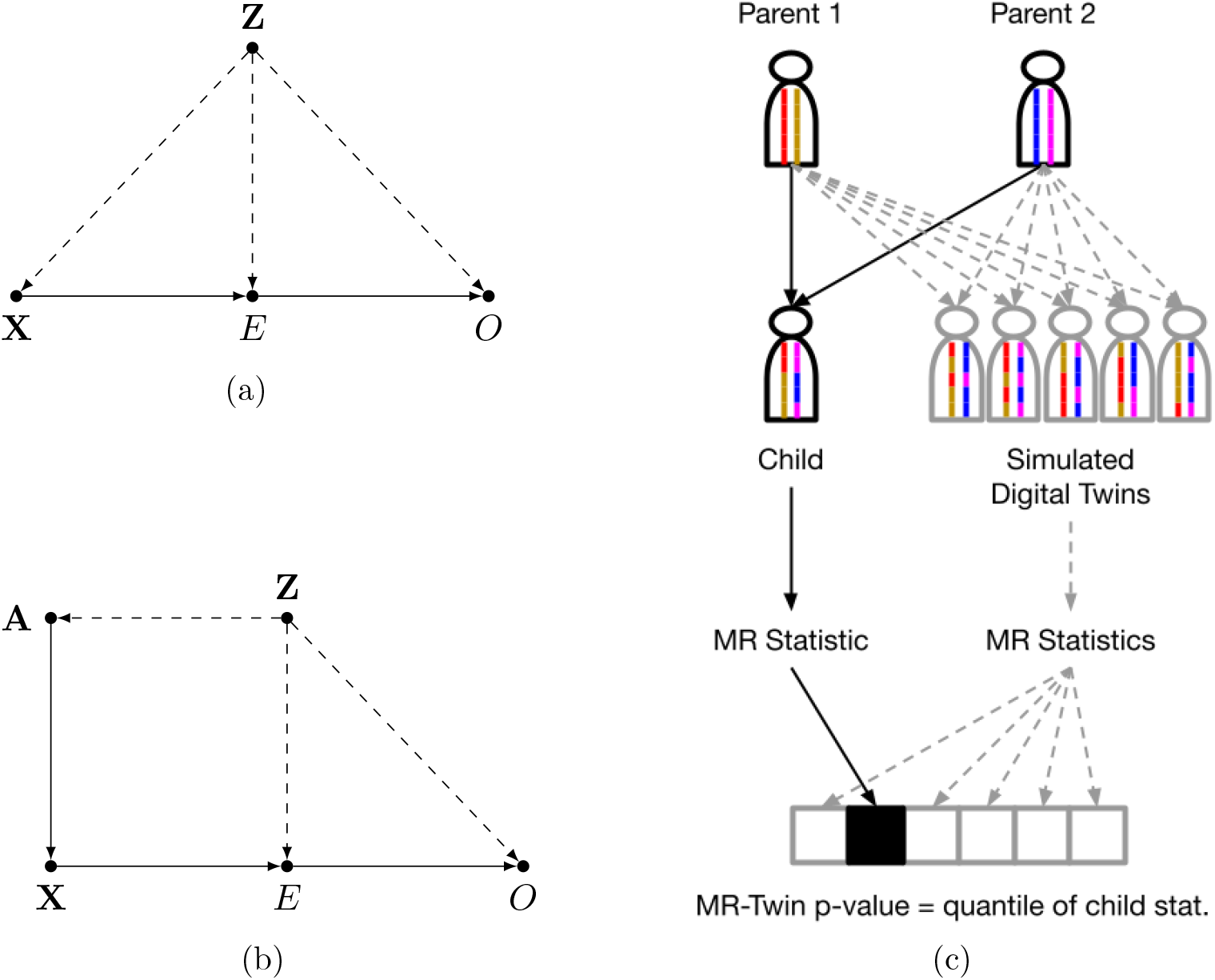
Illustrations of Mendelian Randomization assumptions and the MR-Twin framework. (a) Directed Acyclic Graph (DAG) depicting variables and their relationships in a typical Mendelian Randomization (MR) study, where *X* is the genotypic instrument, *E* is the exposure trait, and. *O* is the outcome trait. An external confounder *Z*, such as population stratification, can cause violations of the MR assumptions. (b) If we have the parental haplotypes *A*, then X is independent of *Z* given *A*. (c) Illustration of the MR-Twin workflow. Digital twin genotypes are sampled from the parental genotypes. MR-Twin is a conditional randomization test, conditioned on *A* and therefore immune to confounding from *Z*, in which the p-value is computed based on the quantile of the true offspring’s MR-Twin statistic compared to the digital twins’ statistics.

MR-Twin is a method that uses family-based genetic data to construct a test for whether the exposure has an effect on the outcome that is immune to confounding from population structure. It is based on the key idea that the genotypes of observed individuals are independent of population structure given the genotypes of the individuals’ parents, since the mechanisms by which genetic information is passed from parents to offspring are known (Figure 1b). In other words, conditioned on the parental genotypes, population structure provides no additional information about the distribution of the offspring’s genotypes. Thus, by conditioning on the parental genotypes, confounding from population stratification can be avoided (Figure 1c), along with confounding from other phenomena such as cross-trait assortative mating and dynastic effects which operate through the parental genotypes (see Figure 1 of Brumpton et al [11]).

We now outline the algorithm in the context of a trio design in which we have genetic data on the parents and the offspring. Let **X** and *O* denote the genotypes and outcome phenotype values respectively for some individual, and let 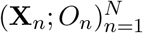, denote these across *N* trios. Also let **P1** and **P2** denote the genotypes of the parents of the individual with genotypes **X**, and let **A** ≔ (**P1, P2**) refer to the set of parental genotypes. Let ***Z*** denote the set of external confounders measured on the same individual, which we define as the set of confounders that satisfy ***X*** ⫫ ***Z***| ***A***. Thus, population stratification is an external confounder (as are assortative mating and dynastic effects) while horizontal pleiotropy is not. The key idea is that we can formulate a hypothesis test of a causal effect conditional on the parental haplotypes ***A***. Bates et al. [18] show that such a test is also a test of the stronger null hypothesis of a causal effect conditional on (***A**, **Z***).

The way that this is accomplished is through a conditional randomization test, similar to the Digital Twin Test proposed by Bates et al in the context of GWAS [18, 19]. The idea is to sample so-called “digital twins” 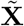 from each set of parents **A** such that 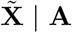 has the same distribution as **X** | **A**, which can easily be accomplished using the laws of Mendelian inheritance (Methods). We construct *B* such random samples across all trios, 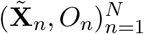, and for each set *b* of twins we compute a test statistic 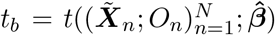 representing the strength of association between the genetically-predicted exposure and the outcome. We also compute a test statistic for the true offspring of the trios, 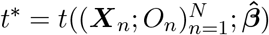.

We can then obtain a p-value for a non-zero causal effect of the exposure on the outcome, 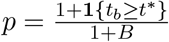. The set of *B* statistics derived from the digital twins represents a null distribution conditioned on the parental genotypes. If there is a true nonzero effect of the exposure on the outcome, we expect the statistic derived from the true offspring to be stronger than statistics derived from digital twins whose genotypes are randomly sampled from the parental genotypes. The test statistic and algorithm are explained in more detail in the Methods section.

### 2.2 MR-Twin controls for arbitrarily strong population stratification confounding in simulations

We compared the performance of MR-Twin to other MR methods via simulations consisting of two populations with allele frequency differences modeled according to the standard Balding-Nichols model [20], following previous works [21, 22, 23, 24, 25, 26]. The procedure for simulating the genotypes is outlined in Algorithm 1. We use this algorithm to simulate “external” samples (non-trio data, e.g. from a biobank), as well as the parents for the trios. The offspring for the trios can then be easily sampled given the parental genotypes (Methods). For each sample, we retain the population label, a binary variable indicating which population each sample belongs to. Unless otherwise specified, each simulation had 50,000 (false positive rate simulations) or 100,000 (power simulations) external samples and 1,000 trio samples evenly split between two populations with fixation index (*F_ST_*)= 0.01, and 100 SNPs, 50 of which were causal for the exposure trait. Unless otherwise specified, the heritability of the exposure trait was set to *h*^2^ = 0.2.

#### Algorithm 1 Simulate genotypes under population structure

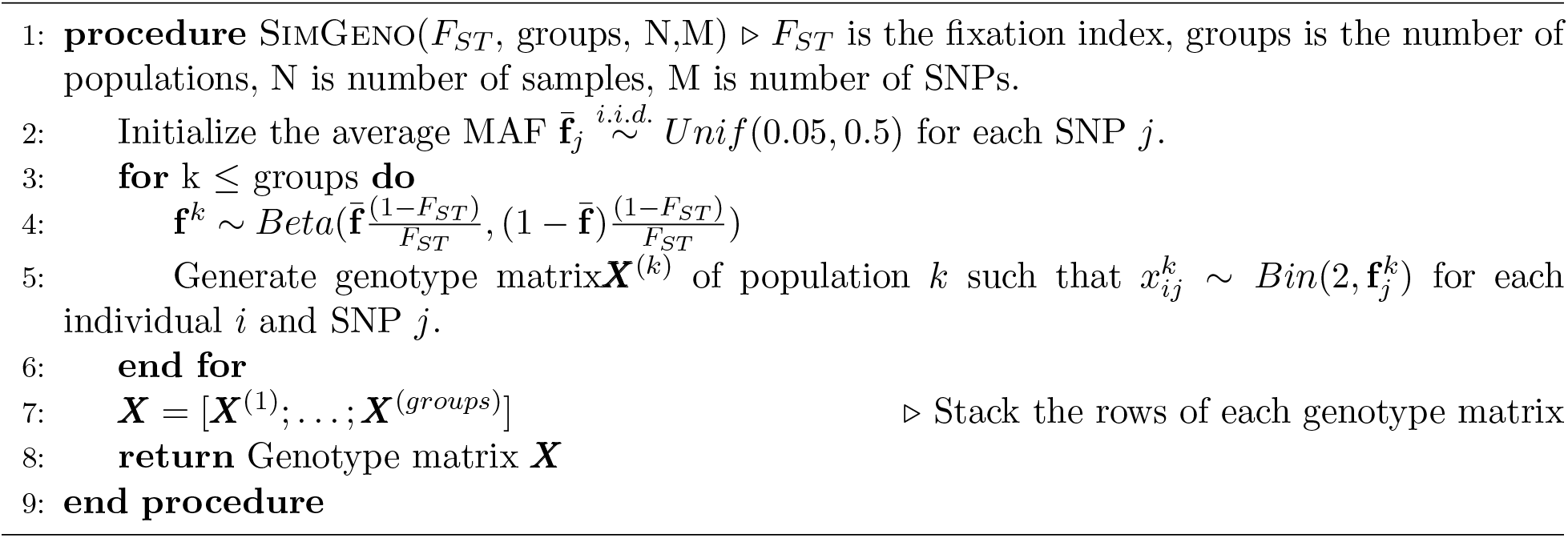

Next, we simulate both the exposure and outcome phenotypes following a linear model, as outlined in Algorithm 2. This model allows the population labels from the first step to have an effect on the exposure and outcome phenotypes, which models population stratification that violates the MR assumptions. We use this setting to assess the false positive rates (FPR) of methods under population stratification, allowing the effects of the population labels on the exposure and outcome phenotypes to range from 0 (no confounding) to 0.8 (substantial confounding). In a separate set of simulations to assess power, we set the confounding effect to 0 and varied the causal effect.

#### Algorithm 2 Simulate population-stratified phenotypes

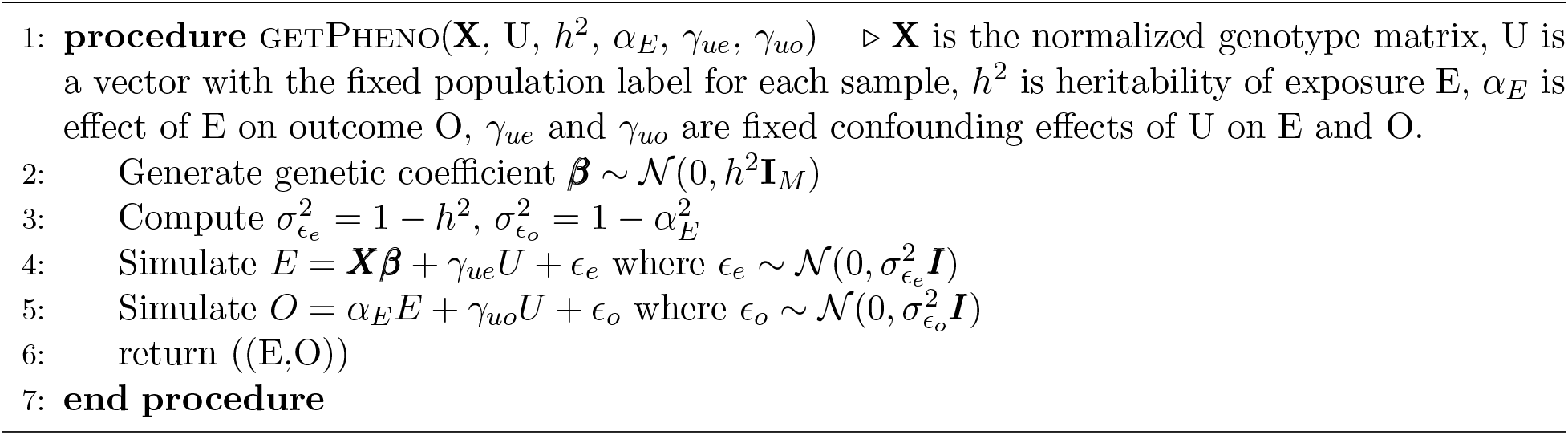

We performed 1000 simulation replicates under these settings, each time simulating a set of external and trio genotypes and phenotypes according to the chosen parameters, performing linear regression between each SNP and the exposure and outcome phenotypes, and using the resulting association statistics as input to each of the MR methods. We excluded SNPs with association p-values of below 0.05/*M* (*M* = 100) with the exposure phenotypes in the external data in order to limit weak instrument bias [8]. The methods we assessed include the trio mode of MR-Twin, standard inverse-variance weighted (IVW) MR [27], MR-Egger [28], the Weighted Median Estimator [29], the Mode-Based Estimator [30], and a method recently introduced by Brumpton et al [11] to use family data to control for confounding due to population stratification and other population-related effects. Brumpton et al provide a suite of methods for different family datasets, following previous work such as [31]; here we focus on the trio-based method they describe [11], and simply refer to that method as “Brumpton” below.

The trio mode of MR-Twin maintained a calibrated FPR irrespective of the strength of confounding (Figure 2a). Non-family-based methods such as IVW, Egger, Median, and Mode all displayed substantially inflated FPR in the face of confounding consistent with their sensitivity to potential residual population stratification. The Brumpton method also displayed slightly inflated FPR, which increased with the strength of the confounding effect, likely due to weak instrument bias [11]. To mitigate the impact of weak instrument bias, we applied a common approach employed in MR studies [8] that involves filtering out variants for which the F-statistic of the association signal is low (*F* < 10 following previous recommendations). This rendered the FPR inflation negligible, but also rendered Brumpton substantially less powerful than MR-Twin, whereas the “unfiltered” mode had similar power to MR-Twin (Figure 2b). Results with confidence intervals are shown in Figure S6. We further investigated the weak instrument bias by running simulations with no SNP filtering based on external data and with increasing numbers of SNPs – settings expected to generate large numbers of weak instruments – and confirmed that Brumpton had greater FPR inflation in these settings while MR-Twin remained calibrated (Figure S7) and did not lose power (Figure S8).

**Figure 2:**
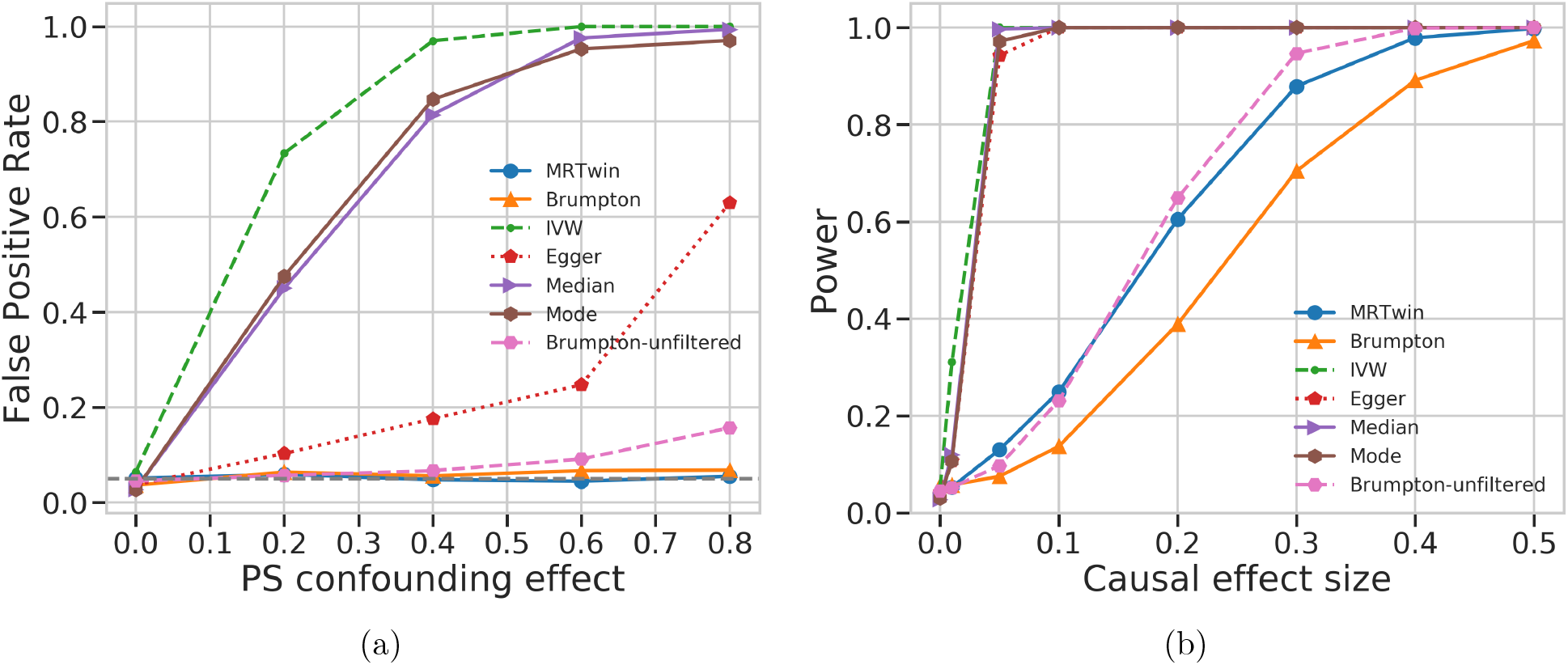
False Positive Rate (FPR) and Power comparison between various methods run on simulated data. (a) False positive rate (y-axis) under varying levels of confounding due to population stratification (PS), with the x-axis describing the magnitude of the confounding effect of population labels on the exposure and outcome trait. (b) Power (y-axis) as a function of the magnitude of the causal effect of the exposure on the outcome trait (x-axis) in a setting with no confounding. Results are averaged over 1000 simulation replicates.

The standard MR methods (IVW, Egger, Median, and Mode), when run on the external data, had substantially higher power than the family-based methods, MR-Twin and Brumpton (Figure 2b). We performed additional simulations to understand if the lower power of MR-Twin was due to the smaller number of trios as opposed to methodological limitations. When applied to the offspring in each trio (Figure 3), the standard MR methods still had substantially inflated FPR (Figure 3a) but similar power to MR-Twin and Brumpton (Figure 3b). We also evaluated the FPR and power of these methods under varying number of trios (Figure S1). We observed that increasing number of trios increased power for all methods, as expected, suggesting that the family-based methods can be expected to obtain increased power as more genetic family data are ascertained in the future. The relative power of the methods remained roughly consistent across these experiments.

**Figure 3:**
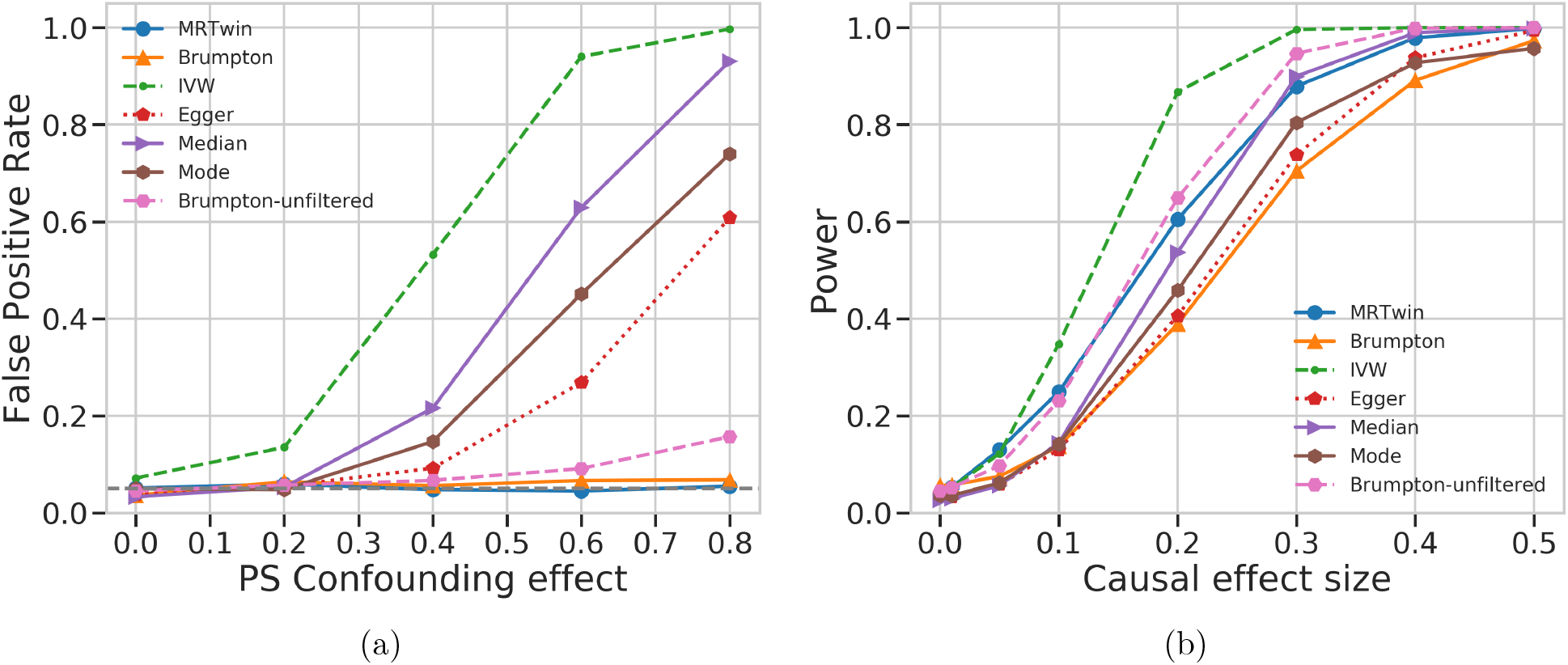
False Positive Rate (FPR) and Power comparison between various methods run on simulated trio data. This is similar to Figure 2 except that IVW, Egger, Median, and Mode are run on the offsprings of the trio dataset instead of the large “external” group of unrelated individuals, such that all methods have the same sample size. (a) False positive rate (y-axis) under varying levels of confounding due to population stratification (PS), with the x-axis describing the magnitude of the effect of the population labels on the exposure and outcome trait. (b) Power (y-axis) as a function of the causal effect size (x-axis). Results are averaged over 1000 simulation replicates.

We performed simulations increasing the magnitude of population structure as measured by the *F_ST_* (without necessarily increasing the confounding strength), and observed that increasing the population structure leads to further FPR inflation for standard MR methods (Figure S2). We observed inflated FPR for standard MR methods even when there is no confounding for large values of *F_ST_* (Figure S2) likely due to correlation or linkage disequilibrium among the genetic variants induced by population structure. The standard implementation of IVW and Egger [32, 33] allows the user to pass in a variant correlation matrix, which removed the FPR inflation with no stratification (Figure S2); other methods such as Median and Mode do not currently have this option.

Next we assessed the runtime of methods run on the trio data (Figure S3). Brumpton, along with non-trio based methods (e.g., IVW), had similar run times (<1-5 seconds per simulation replicate); for succinctness, only Brumpton is shown. MR-Twin (with 100 simulated digital twins) took roughly one minute per simulation replicate under the simulation settings described above, with time increasing to up to four minutes if the number of families or SNPs was increased. The number of digital twins to simulate for MR-Twin involves a trade-off between speed and stability of results. We assessed the stability of MR-Twin with different numbers of digital twins, with results shown in Figure S5. We interpreted these findings as indicating that 100 digital twins are likely stable enough for simulations for which many replicates are run and speed is a priority, but 1000 or more digital twins is recommended for one-off real data analysis. Therefore we simulated 100 digital twins in our simulations and 1000 in our real data analysis. We note, however, that while the MR-Twin runtime increases linearly with the number of digital twins simulated, the generation and statistic computation for the digital twins can be done in parallel, so many twins can be simulated efficiently given multiple compute cores or nodes. For clarity of results, we did not take advantage of this in our runtime assessment.

Finally, MR-Twin also enables users to use parent-child duo or sibling datasets (Methods). We assessed the performance of these modes versus the trio mode of MR-Twin (Figure S4). We found that the duo and sibling modes, while having lower FPR than most standard MR methods, did not maintain a calibrated FPR at high levels of confounding, which is expected since the precise sampling of offspring genotypes from parents is not possible when either or both of the parental genotypes are not available.

### 2.3 Application to trio data in the UK Biobank

In order to assess the results given by MR-Twin relative to other approaches in a real data context, we next applied MR-Twin and four other MR methods (IVW, Egger, Median, and the Brumpton et al. method [11]) to 144 real trait pairs in the UK Biobank [1]. These consisted of all pairwise combinations of 12 metabolic, anthropometric, and socioeconomic traits that were widely measured among the UK Biobank participants. We isolated 955 White British genetic trios from the full UK Biobank dataset (Supplementary Note) and used PLINK [34] to run linear regression on the remaining unrelated White British individuals for these 12 traits, including the top 20 principal components (PCs), age, and sex as covariates. The genetic instruments selected for each analysis were the SNPs with genome-wide significant p-values (< 5.0 × 10^-8^) for the exposure trait, after linkage disequilibrium (LD) pruning was performed so that none of these instruments were in substantial LD with one another (Supplementary Note). Ignoring the degenerate cases where the exposure and outcome were the same trait or where there were no significant SNPs for the exposure trait (as was the case for the Townsend Deprivation Index [Deprivation]), there were 121 usable trait pairs.

Table 1 shows the results for six selected trait pairs (excluding Median for brevity because it gave similar results to IVW), while Supplementary Table S2 shows the full set of analyses. (Brumpton was run with several different variant filtering settings to assess the impact of potential weak instrument bias (Supplementary Note); results for all runs are given in Supplementary Table S2.) For Table 1, we selected six analyses: two positive controls representing causal effects that are true by definition (LDL Cholesterol → Total Cholesterol and Weight → Body Mass Index [BMI]), two negative controls that represent seemingly implausible effects (Glucose → Deprivation and Height → Body Fat), and two trait pairs with unclear or conflicting evidence (BMI → Diastolic Blood Pressure [DBP] and BMI → Deprivation). In particular, previous studies have identified a significant effect for BMI → DBP [4] and for BMI → Deprivation in women [35] with IVW analysis, although Egger analysis did not replicate the significant findings in either case [4, 35].

**Table 1:** Traditional MR results and MR-Twin results on selected trait pairs from the UK Biobank. Bold numbers are significant at *p* < 0.05. Note that 9.99 × 10^-4^ = 1/1001 is the minimum p-value for MR-Twin in this expriment, as 1000 digital twins were generated. Chol. = Cholesterol; BMI = Body Mass Index; DBP = Diastolic Blood Pressure; Deprivation = Townsend Deprivation Index.

| Traits | MR P-Values |  |  | MR-Twin P-Value |
| --- | --- | --- | --- | --- |
|  | IVW | Egger | Brumpton | MR-Twin |
| LDL Chol. → Total Chol. | $< \mathbf{10^{-300}}$ | $< \mathbf{10^{-300}}$ | $\mathbf{1.64} \times \mathbf{10^{-11}}$ | $\leq \mathbf{9.99} \times \mathbf{10^{-4}}$ |
| Weight → BMI | $< \mathbf{10^{-300}}$ | $< \mathbf{10^{-300}}$ | $\mathbf{4.80} \times \mathbf{10^{-6}}$ | $\leq \mathbf{9.99} \times \mathbf{10^{-4}}$ |
| BMI → DBP | $\mathbf{2.24} \times \mathbf{10^{-26}}$ | $5.64 \times 10^{-1}$ | $\mathbf{3.46} \times \mathbf{10^{-2}}$ | $2.69 \times 10^{-1}$ |
| BMI → Deprivation | $\mathbf{1.18} \times \mathbf{10^{-19}}$ | $\mathbf{7.53} \times \mathbf{10^{-3}}$ | $9.99 \times 10^{-2}$ | $8.79 \times 10^{-2}$ |
| Glucose → Deprivation | $1.54 \times 10^{-1}$ | $2.09 \times 10^{-1}$ | $6.61 \times 10^{-1}$ | $1.91 \times 10^{-1}$ |
| Height → Body Fat | $9.55 \times 10^{-1}$ | $9.83 \times 10^{-2}$ | $6.73 \times 10^{-1}$ | $5.09 \times 10^{-1}$ |

All methods performed as expected on the controls, with highly significant p-values for positive controls and insignificant p-values for negative controls. For BMI → DBP, IVW and Brumpton yielded significant results while Egger and MR-Twin did not. For BMI → Deprivation, IVW and Egger yielded significant results while Brumpton and MR-Twin did not. In general, IVW tended to yield much stronger p-values than other methods and the family-based methods (Brumpton and MR-Twin) tended to be conservative (Supplementary Table S2), in line with our simulation results. In particular, of the 121 usable trait pairs, IVW identified 78 as significant, Egger identified 56 as significant, Brumpton identified 20 as significant, and MR-Twin identified 19 as significant.

## 3 Discussion

We introduced MR-Twin, a method for testing causal effects between pairs of traits within a Mendelian Randomization (MR) framework, which is provably robust to confounding of any strength resulting from population stratification. We demonstrated that existing MR methods, including those designed to correct for confounding resulting from horizontal pleiotropy, are prone to false positives when there is confounding from population stratification. We further demonstrated that MR-Twin can leverage parent-child duo or sibling data if trio data is not available, although FPR is not fully controlled in those cases.

While population stratification was the focus of this paper, the MR-Twin framework also provides immunity to several other types of confounding effects. Theory dictates that MR-Twin is immune to confounding from familial effects such as assortative mating and dynastic effects since these effects operate through the parental genotypes (see Figure 1 of Brumpton et al [11]), though we do not explicitly test this in this manuscript. As recently demonstrated, cross-trait assortative mating is pervasive and impacts many common genetic analyses [16] including MR [15], so this represents another valuable aspect of MR-Twin even if population stratification is believed to be well-controlled in a particular study. In general, MR-Twin is immune to any confounder that is independent of the genotypes of offspring given the genotypes of their parents.

In addition to population and familial effects, we highlight two under-appreciated sources of bias in MR studies, both of which MR-Twin avoids without requiring the user to modify any parameters or arguments. The first is weak instrument bias [8], which can bias the effect estimate of standard MR methods, including the Brumpton approach [11]. This accounts for the Brumpton method yielding inflated FPRs when the confounding effects were strong (Figure 2a). One of the most common ways to control for weak instrument bias is by filtering out variants with a weak association signal, often with a threshold of *F* < 10 for the association between a variant and the exposure trait. However, this procedure has been criticized [8] and may not fully correct for weak instrument bias. Other MR methods may also be affected by this bias. In two-sample study designs, the direction of the bias is towards the null rather than the confounded exposure-outcome association estimate [36], but the bias remains.

Additionally, we found that standard MR methods (IVW, Egger, Median, Mode, etc) may have inflated FPR even when there is no population stratification when there is population structure in the data, with the inflation scaling with increased fixation index (Figure S2). The reason for this is that, even though the variants were simulated independently, they were correlated with one another through the population labels. For example, suppose we have two variants, *X*1 and *X*2 and a population label *U*. The causal diagram for these three variables is *X*1 ← *U* → *X*2, so *X*1 and *X*2 are correlated. This is both a point of caution for performing MR simulations and a practical concern for users, for whom a SNP correlation matrix may not always be available or computable if they are working with summary data.

MR-Twin avoids both of these issues, without requiring the user to specify a SNP correlation matrix or apply various approaches to mitigate weak instrument bias. First, both MR-Twin and Brumpton avoid the correlated-variant issue because they condition on parental genotypes, severing the link between the offspring genotypes and the population structure. Second, MR-Twin would not lose FPR calibration due to weak instrument bias, because this phenomenon has nothing to do with the aspects of the MR-Twin test that guarantee immunity from confounding due to population and familial effects (sampling digital twin genotypes conditioned on parental genotypes). Theoretically, it is possible that the bias in the MR effect estimate used in the MR-Twin statistic (Methods) could lower power, but because the MR effect estimate equally affects both the digital twin statistics and the true offspring statistics, a reduction in power seems unlikely and was not observed empirically (Figure S8).

There is extensive literature on family-based methods for avoiding confounding due to population structure in genome-wide association studies or linkage analysis [37, 38, 39, 40, 31, 41, 42]. One prominent example is the transmission disequilibrium test (TDT) [42] and the more-recent polygenic TDT (pTDT) [37]. Bates et al [18] compare the digital twin test (DTT) to the TDT and show that the DTT is a generalization of the TDT and highlight some of its benefits. Because it is not immediately obvious how to adapt the TDT and pTDT to MR, we do not evaluate their potential use in this context.

There are several considerations that come into play when applying the MR-Twin method, which we note here. First, the number of digital twins simulated involves a trade-off between speed and precision (Figure S5). While MR-Twin was slower than competing MR methods (Figure S3), it still ran in a few minutes or less per run on both simulated and real data analyses, justifying the use of a fairly large number of digital twins if possible. Consequently, we recommend 1000 or more digital twins for real data analysis, which should be computationally feasible and precise (and, again, parallelization can make this quite efficient). 100 digital twins is likely sufficient in simulations where there are many replicates and speed is the paramount concern. Second, the populations of the external and family datasets should be similar. This is natural for biobanks like the UK Biobank, but can be more challenging when attempting to combine separate datasets. Third, care should be taken to ensure that the normalization method used and covariates controlled-for are similar in the external and trio datasets in order to avoid potential loss of power.

While the genetic trio offspring used in our UK Biobank analysis were all adults (as all participants in this dataset were aged 40-69 at collection time [1]), other trio datasets may contain young children. This is a potential issue because some commonly-analyzed traits such as height and weight may not have the same relationship in youths or adolescents as they do in adults, and variants that affect these traits may not yet have realized their full effect in the children yet. Dealing with such time-varying exposures in the context of mendelian randomization is an area of ongoing research [43], and it is not clear how this would impact MR-Twin results. Even when the offspring of the trios are all adults, it may be difficult to adequately sample certain traits. For example, we were not able to perform MR analysis for complex traits such as heart disease, since none of the offspring in our sample had developed heart disease, largely because all offspring in our sample were aged 40-49.

Several extensions to the methods presented here are also possible. While we explored continuous traits in this paper, further work needs to be done to apply MR-Twin to binary phenotypes such as disease labels. First, a different statistic such as binary cross entropy (rather than our negative squared loss statistic) may be more appropriate. Second, the use of the external effect size estimates in the statistic may have to be modified, depending on the regression method used and the interpretation of the estimates. For example, it would be inappropriate to replace the effect size estimates in our statistic with odds ratios produced by logistic regression. Even for linear data, it is possible that a different statistic than the one we proposed would be more powerful in some situations. Finding most-powerful statistics for a given significance threshold is a direction for future work.

In the Digital Twin Test paper [18], Bates et al propose using a Hidden Markov Model (HMM) to simulate digital twins from the parental haplotypes, the latter being generated by phasing the parental genotypes. For the simplicity of avoiding this phasing step and due to the fact that genetic instruments in MR studies are usually selected to be roughly independent [27], we used a simpler method for simulating digital twins using binomial draws from the parental genotypes (Methods). However, the variants used may not be independent even if they appear to be [27], or one may wish to include correlated variants to increase power. Extending MR-Twin to perform the HMM-based digital twin simulation could therefore increase power.

## 4 Methods

### 4.1 The MR-Twin Framework

We first introduce the standard Mendelian Randomization (MR) model, without any confounding. Suppose that for a collection of *N* individuals we obtain their genotypes at M SNPs, and a phenotypic measure for an exposure trait and an outcome trait. For a given individual *n* we denote the genotype vector as **X**_*n*_, the genotype at some SNP *j* as **X**_*nj*_, the exposure trait as *E_n_*, and the outcome trait as *O_n_*. Let 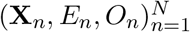 denote the collection of these genotypes and traits over all *N* individuals, where (**X**_*n*_) is an (*N* × *M*) matrix and (*E_n_*) and (*O_n_*) are (*N* × 1) vectors. Finally, let **X**, *E*, and *O* refer to the genotype vector, exposure trait, and outcome trait for a generic individual.

MR uses the genetic “instruments” **X** to estimate the effect of an “exposure” trait *E* on an “outcome” trait *O*. This estimate is valid regardless of any confounder **U** of the association between *E* and *O*, assuming that the following conditions hold [7]:

1. The genetic instrument **X** is significantly associated with the exposure trait *E*;
2. The genetic instrument **X** is independent of any variables (such as those in **U**) that confound the relationship between *E* and the outcome trait *O*;
3. The genetic instrument **X** is not associated with *O* except due to its association with *E*.

The latter two criteria can be captured by the independence statement

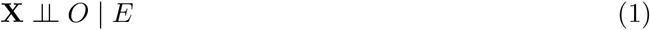

Assuming these conditions hold, and assuming a linear model for the relationships between the genotypes and phenotypes (a typical assumption in MR analyses), we can test the null hypothesis that there is no direct causal effect of *E* on *O*,

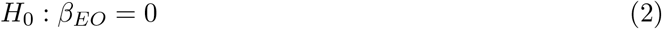

where *β_EO_* is not obtained by direct regression but rather via instrumental variables estimators such as the ratio estimator *β_EO_* = *β*_*XO*_/*β_XE_* (when a single instrument is used) or by two stage least squares or inverse-variance weighting (when multiple instruments are used) [27].

However, in the case where we have residual population stratification, denoted **Z** (Figure 1a), this independence assumption is violated. This is because, using terminology from Pearl’s graphical formalism [44], **X** ← **Z** → *O* is a fork and thus there is a backdoor path between **X** and *O*, so the two are not marginally independent. Conditioning on *E* fails to block this backdoor path. Residual population stratification generally cannot be controlled for directly, though approaches such as Principal Components Analysis (PCA) and Linear Mixed Models (LMMs) have been used to reduce its effect [9].

MR-Twin (Figure 1c) is a method that uses family-based genetic data to avoid this confounding. Suppose that, corresponding to each individual’s genotypes **X**, we also observe the genotypes **P1** and **P2** of their parents (we later relax the trio assumption to allow for parent-child duo or sibling data). Let **A** ≔ (**P1**, **P2**). According to the graphical criteria for d-separation developed by Pearl [44], **A** d-separates **X** from **Z** (Figure 1b):

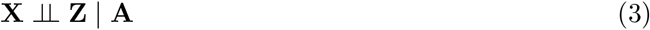

This means that, assuming **X** does not affect some unmeasured variable which in turn affects *O* (i.e. no horizontal pleiotropy),

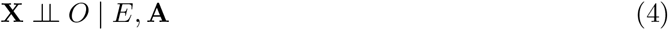

thereby satisfying the MR conditions regardless of any residual population stratification.

As shown by Bates et al [18], the Digital Twin Test framework outlined in Algorithm 3 can be used to perform a hypothesis test conditioned on **A**. The resulting test involves computing the test statistic 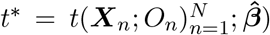 (we give the statistic used in this paper in “MR-Twin Test Statistic Incorporating External Weights” below). To perform a test, we construct *B* random samples 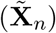 where each 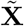 is a random sample given **A** with the same distribution as **X** given **A** (such a sample can be easily constructed using Mendelian inheritance; see “Generating Digital Twins” below). We refer to these samples as “digital twins”. For each such sample *b*, we then compute 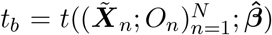, representing a null distribution of genotypes conditioned on the parental genotypes. This, in turn, gives us a p-value for 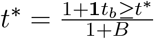, where *B* is the total number of permutations we perform. The MR-Twin test is therefore a kind of conditional randomization test [18, 19].

#### Algorithm 3 MR-Twin

1. Input: Effect sizes for SNPs: 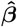, trio data 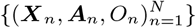
2. Compute the MR-Twin test statistic 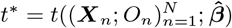
3. For b = 1 to B:

a. Sample digital twins 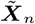 given their ancestors ***A**_n_*.
b. Compute the MR-Twin test statistic 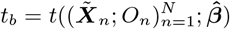
4. 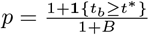

Output: p-value: *p*

Importantly, the proposed algorithm can leverage effect size estimates 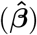 from any external GWAS datasets (even GWAS datasets where such estimates might be biased due to population stratification) while providing valid tests. The proposed algorithm is robust to any external con-founder satisfying Equation 3, such as population stratification, assortative mating, and dynastic effects.

Next, we detail the MR-Twin test statistic, digital twin generation algorithms, and formal proofs of the exchangeability of digital twins with each other and their real counterparts.

### 4.2 Conditional Randomization Test for Mendelian Randomization

The MR-Twin test is related to the digital twin test [18] and likewise is a kind of conditional randomization test [19]. Like the digital twin test, MR-Twin leverages the fact that offspring genotypes are conditionally independent of “external” confounders such as population structure given the parental genotypes and uses a conditional randomization test to test the weaker, but equivalent, null hypothesis of no effect conditioned upon the parental genotypes.

Let **X** be a vector of offspring genotypes, and let **A** be the genotype vectors of the two parents of the offspring. **A** may be directly observed, as in trio data, or inferred using parent-child duo or sibling data (see “Generating Digital Twins”). Let **Z** be one or more “external” confounders, defined [18] as

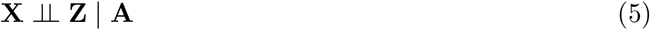

Thus, population structure is an external confounder, while horizontal pleiotropic traits are not.

We therefore have

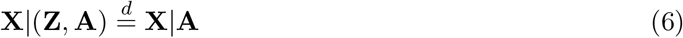

Assuming that all confounders are external and that **X** is significantly associated with *E*, *O* is independent of **X** given **A** under the MR null hypothesis that *E* has no effect on *O*. This is because **X** would not have any effects on *O* mediated by *E* (since *E* does not affect *O* under the MR null hypothesis), and all paths not through *E* are blocked by conditioning on **A** as shown in Equation 6. We therefore want to test

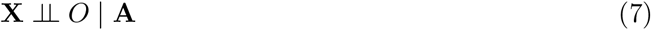

If this holds, then we cannot rule out that either **X** has no effect on *E* or *E* has no effect on *O*. We test this null hypothesis via a conditional randomization test [19].

In testing this null hypothesis, it is helpful to be able to leverage SNP effect sizes estimated from large, external datasets (such as publicly released summary statistics for resources like the UK Biobank [1]), as this will often yield more statistically significant variants and better effect size estimates than those generated using small genetic family datasets. We therefore note that the following property also holds:

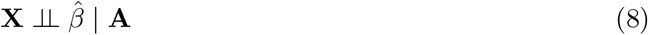

where we use the shorthand 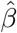 to refer to the estimated effect sizes of each SNP on the exposure and outcome traits.

We construct “digital twins” 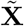 sampled from the parental genotypes via Mendelian inheritance (see “Generating Digital Twins”) such that

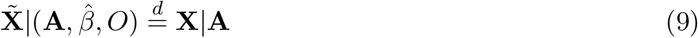

Given equations 7 and 8, we have the following under the null hypothesis:

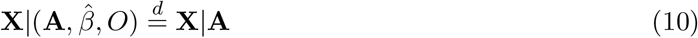

It follows from equations 9 and 10 that the digital twins are exchangeable under the null hypothesis:

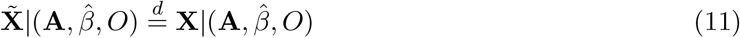

Therefore, given some statistic 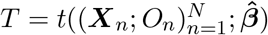, where *N* is the number of families,

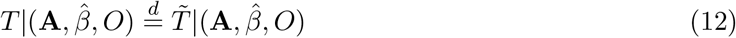

under the null, where 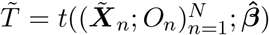. We can then use the procedure outline in Algorithm 3 to obtain a p-value for this test statistic [19].

### 4.3 MR-Twin Test Statistic Incorporating External Weights

We construct a test statistic based on a negative sum of squares loss when using **X** to predict *O* via an MR estimate for the effect of *E* on *O*. First, we leverage the effect sizes from the external dataset of the genotype on the exposure trait 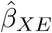 to obtain the genetically-predicted exposure trait values:

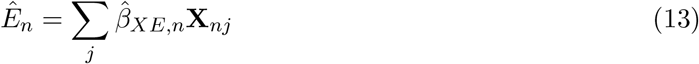

for each individual *n* and SNP *j*. We then compute the MR estimate for the effect of the exposure trait on the outcome trait, 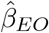. This estimate may be a conventional Inverse Variance Weighted (IVW) estimate [27] or various statistics designed to be robust to pleiotropy such as the Egger-based statistic [28], the weighted median statistic [29], or others. We then predict the outcome trait for each individual *n* as 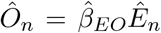. Finally, we compute the negative squared error of these predictions 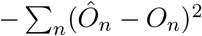. The full statistic is then

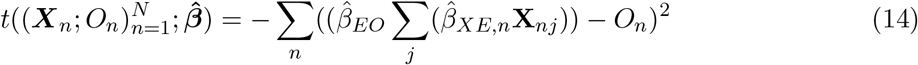

### 4.4 Generating Digital Twins

We have assumed that trio data is available thus far for simplicity. However, the MR-Twin framework can also be used when parent-child duo data or sibling data are available. Here we discuss the algorithms used to generate digital twins given trio, parent-child duo, or sibling data.

#### 4.4.1 Trio and Duo modes

We assume that the SNPs used in the MR instrument are independent, a common assumption when multi-SNP instruments are used in MR [27]. Therefore, we separately sample the genotype of each SNP of the digital twin given the parent and/or offspring genotypes at that SNP. Let (**D**_*n*_) be the (*N* × *M*) matrix of digital twin genotypes we will sample, corresponding to the true “offspring” genotypes in (**X**_*n*_). Further, let n index some family and *j* index some SNP, such that **P1**_*nj*_ (for example) is the genotype for one parent in family *n* at SNP *j*. If we have both parents available, sampling **D**_*nj*_ is straightforward. Because the SNPs are considered independent, we do not need to know the parental haplotypes. If a parental genotype **P1**_*nj*_ is 0 or 2, respectively, then a 0 or 1, respectively, is inherited by **D**_*nj*_. If the parent genotype is 1, then either 0 or 1 is inherited with 50% probability each. **D**_*nj*_ inherits alleles from the two parents independently. This can be summarized as

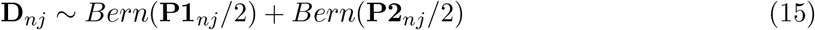

where *Bern* stands for the Bernoulli distribution, for each family *n* and SNP *j*.

If we only have one parent genotype available, then following Bates et al [18], we fix the offspring’s haplotype from the unobserved parent and only simulate a random draw from the observed parent’s haplotype. If the observed parent is homozygous, then the allele inherited from that parent is fixed as well, so **D**_*nj*_ = **X**_*nj*_. Otherwise, the allele inherited from this parent will be *Bern*(0.5). Similarly to the above, the model for the allele from the other parent can be written as *Bern*(**X**_*nj*_/2). Thus, if the parent is a heterozygote, we have

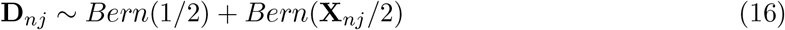

#### 4.4.2 Sibling mode

In the case where we observe sibling genotypes but not the genotypes of their parents, we assessed two potential approaches. In either case, the observed sibling information is used to infer the probabilities of digital twin genotypes based on the fact that the sibling genotypes give information about the probabilities of various parental genotypes. For instance, a child with a 2 genotype at a SNP guarantees that neither parent has a 0 genotype at that SNP, and makes it more likely that the parents have 2 genotypes than 1 genotypes. Most simply, if one sibling has a 2 genotype at a SNP and the other sibling has a 0, then the parents must both be heterozygotes. In all other cases, approximation is needed.

The first approach is straightforward and involves randomly drawing two haplotypes from the observed sibling haplotypes to generate a digital twin. This shuffling approach gives a rough approximation of the likelihood of digital twin genotypes given the information the observed siblings provide. The second approach, described in the Supplementary Note, involves using the sibling data to infer a distribution over the possible parents, then performing a weighted random draw of digital siblings based on those parents. In practice, we found that the shuffling approach was faster and yielded lower FPR than the probabilistic approach while achieving similar power, so we used the shuffling approach for the results in this paper.

## 5 Code and Data Availability

The code implementing the MR-Twin package can be found at: https://github.com/nlapier2/MR-Twin. Scripts and instructions for repeating the experiments in this paper can be found at: https://github.com/nlapier2/MRTwin-replication. Please note that UK Biobank genotypes are not publicly released, so those wishing to replicate the experiments will first have to get access to that data via https://www.ukbiobank.ac.uk/enable-your-research/apply-for-access.

## 6 Acknowledgements

This research has been conducted using the UK Biobank Resource under application number 33127. B.F. and S.S were supported in part by NIH R35GM125055 and NSF CAREER-1943497, III2106908.

## References

[1] C. Bycroft, C. Freeman, D. Petkova, G. Band, L. T. Elliott, K. Sharp, A. Motyer, D. Vukcevic, O. Delaneau, J. O’Connell, et al., “The uk biobank resource with deep phenotyping and genomic data,” Nature, vol. 562, no. 7726, pp. 203–209, 2018.

[2] G. Hemani, J. Zheng, B. Elsworth, K. H. Wade, V. Haberland, D. Baird, C. Laurin, S. Burgess, J. Bowden, R. Langdon, et al., “The mr-base platform supports systematic causal inference across the human phenome,” elife, vol. 7, p. e34408, 2018.

[3] K. H. Wade, D. Carslake, N. Sattar, G. Davey Smith, and N. J. Timpson, “Bmi and mortality in uk biobank: revised estimates using mendelian randomization,” Obesity, vol. 26, no. 11, pp. 1796–1806, 2018.

[4] D. M. Lyall, C. Celis-Morales, J. Ward, S. Iliodromiti, J. J. Anderson, J. M. Gill, D. J. Smith, U. E. Ntuk, D. F. Mackay, M. V. Holmes, et al., “Association of body mass index with cardiometabolic disease in the uk biobank: a mendelian randomization study,” JAMA cardiology, vol. 2, no. 8, pp. 882–889, 2017.

[5] P. C. Haycock, S. Burgess, A. Nounu, J. Zheng, G. N. Okoli, J. Bowden, K. H. Wade, N. J. Timpson, D. M. Evans, P. Willeit, et al., “Association between telomere length and risk of cancer and non-neoplastic diseases: a mendelian randomization study,” JAMA oncology, vol. 3, no. 5, pp. 636–651, 2017.

[6] C. L. Haase, A. Tybjærg-Hansen, A. Ali Qayyum, J. Schou, B. G. Nordestgaard, and R. Frikke-Schmidt, “Lcat, hdl cholesterol and ischemic cardiovascular disease: a mendelian randomization study of hdl cholesterol in 54,500 individuals,” The Journal of Clinical Endocrinology & Metabolism, vol. 97, no. 2, pp. E248–E256, 2012.

[7] D. A. Lawlor, R. M. Harbord, J. A. Sterne, N. Timpson, and G. Davey Smith, “Mendelian randomization: using genes as instruments for making causal inferences in epidemiology,” Statistics in medicine, vol. 27, no. 8, pp. 1133–1163, 2008.

[8] S. Burgess, S. G. Thompson, and C. C. G. Collaboration, “Avoiding bias from weak instruments in mendelian randomization studies,” International journal of epidemiology, vol. 40, no. 3, pp. 755–764, 2011.

[9] A. L. Price, N. A. Zaitlen, D. Reich, and N. Patterson, “New approaches to population stratification in genome-wide association studies,” Nature reviews genetics, vol. 11, no. 7, pp. 459–463, 2010.

[10] J. P. Cook, A. Mahajan, and A. P. Morris, “Fine-scale population structure in the uk biobank: implications for genome-wide association studies,” Human Molecular Genetics, vol. 29, no. 16, pp. 2803–2811, 2020.

[11] B. Brumpton, E. Sanderson, K. Heilbron, F. P. Hartwig, S. Harrison, G. Å. Vie, Y. Cho, L. D. Howe, A. Hughes, D. I. Boomsma, et al., “Avoiding dynastic, assortative mating, and population stratification biases in mendelian randomization through within-family analyses,” Nature communications, vol. 11, no. 1, pp. 1–13, 2020.

[12] J. J. Berg, A. Harpak, N. Sinnott-Armstrong, A. M. Joergensen, H. Mostafavi, Y. Field, E. A. Boyle, X. Zhang, F. Racimo, J. K. Pritchard, et al., “Reduced signal for polygenic adaptation of height in uk biobank,” Elife, vol. 8, p. e39725, 2019.

[13] S. Haworth, R. Mitchell, L. Corbin, K. H. Wade, T. Dudding, A. Budu-Aggrey, D. Carslake, G. Hemani, L. Paternoster, G. D. Smith, et al., “Apparent latent structure within the uk biobank sample has implications for epidemiological analysis,” Nature communications, vol. 10, no. 1, pp. 1–9, 2019.

[14] C. Cinelli, N. LaPierre, B. Hill, S. Sankararaman, and E. Eskin, “Robust mendelian randomization in the presence of residual population stratification, batch effects and horizontal pleiotropy,” bioRxiv, 2020.

[15] F. P. Hartwig, N. M. Davies, and G. Davey Smith, “Bias in mendelian randomization due to assortative mating,” Genetic epidemiology, vol. 42, no. 7, pp. 608–620, 2018.

[16] R. Border, G. Athanasiadis, A. Buil, A. J. Schork, N. Cai, A. I. Young, T. Werge, J. Flint, K. S. Kendler, S. Sankararaman, et al., “Cross-trait assortative mating is widespread and inflates genetic correlation estimates,” Science, vol. 378, no. 6621, pp. 754–761, 2022.

[17] J.-B. Pingault, P. F. O’reilly, T. Schoeler, G. B. Ploubidis, F. Rijsdijk, and F. Dudbridge, “Using genetic data to strengthen causal inference in observational research,” Nature Reviews Genetics, vol. 19, no. 9, pp. 566–580, 2018.

[18] S. Bates, M. Sesia, C. Sabatti, and E. Candès, “Causal inference in genetic trio studies,” Proceedings of the National Academy of Sciences, vol. 117, no. 39, pp. 24117–24126, 2020.

[19] E. Candes, Y. Fan, L. Janson, and J. Lv, “Panning for gold:’model-x’knockoffs for high dimensional controlled variable selection,” Journal of the Royal Statistical Society: Series B (Statistical Methodology), vol. 80, no. 3, pp. 551–577, 2018.

[20] D. J. Balding and R. A. Nichols, “A method for quantifying differentiation between populations at multi-allelic loci and its implications for investigating identity and paternity,” Genetica, vol. 96, no. 1, pp. 3–12, 1995.

[21] A. Ochoa and J. D. Storey, “Estimating fst and kinship for arbitrary population structures,” PLoS genetics, vol. 17, no. 1, p. e1009241, 2021.

[22] M. P. Conomos, A. P. Reiner, B. S. Weir, and T. A. Thornton, “Model-free estimation of recent genetic relatedness,” The American Journal of Human Genetics, vol. 98, no. 1, pp. 127–148, 2016.

[23] G. Chen, A. Yuan, D. Shriner, F. Tekola-Ayele, J. Zhou, A. R. Bentley, Y. Zhou, C. Wang, M. J. Newport, A. Adeyemo, et al., “An improved fst estimator,” Plos one, vol. 10, no. 8, p. e0135368, 2015.

[24] M. J. Hubisz, D. Falush, M. Stephens, and J. K. Pritchard, “Inferring weak population structure with the assistance of sample group information,” Molecular ecology resources, vol. 9, no. 5, pp.1322–1332, 2009.

[25] A. L. Price, N. J. Patterson, R. M. Plenge, M. E. Weinblatt, N. A. Shadick, and D. Reich, “Principal components analysis corrects for stratification in genome-wide association studies,” Nature genetics, vol. 38, no. 8, pp. 904–909, 2006.

[26] J. K. Pritchard, M. Stephens, and P. Donnelly, “Inference of population structure using multilocus genotype data,” Genetics, vol. 155, no. 2, pp. 945–959, 2000.

[27] S. Burgess, A. Butterworth, and S. G. Thompson, “Mendelian randomization analysis with multiple genetic variants using summarized data,” Genetic epidemiology, vol. 37, no. 7, pp. 658–665, 2013.

[28] J. Bowden, G. Davey Smith, and S. Burgess, “Mendelian randomization with invalid instruments: effect estimation and bias detection through egger regression,” International journal of epidemiology, vol. 44, no. 2, pp. 512–525, 2015.

[29] J. Bowden, G. Davey Smith, P. C. Haycock, and S. Burgess, “Consistent estimation in mendelian randomization with some invalid instruments using a weighted median estimator,” Genetic epidemiology, vol. 40, no. 4, pp. 304–314, 2016.

[30] F. P. Hartwig, G. Davey Smith, and J. Bowden, “Robust inference in summary data mendelian randomization via the zero modal pleiotropy assumption,” International journal of epidemiology, vol. 46, no. 6, pp. 1985–1998, 2017.

[31] D. Fulker, S. Cherny, P. Sham, and J. Hewitt, “Combined linkage and association sib-pair analysis for quantitative traits,” The American Journal of Human Genetics, vol. 64, no. 1, pp. 259–267, 1999.

[32] O. O. Yavorska and S. Burgess, “Mendelianrandomization: an r package for performing mendelian randomization analyses using summarized data,” International journal of epidemiology, vol. 46, no. 6, pp. 1734–1739, 2017.

[33] J. R. Broadbent, C. N. Foley, A. J. Grant, A. M. Mason, J. R. Staley, and S. Burgess, “Mendelianrandomization v0. 5.0: updates to an r package for performing mendelian randomization analyses using summarized data,” Wellcome Open Research, vol. 5, 2020.

[34] S. Purcell, B. Neale, K. Todd-Brown, L. Thomas, M. A. Ferreira, D. Bender, J. Maller, P. Sklar, P. I. De Bakker, M. J. Daly, et al., “Plink: a tool set for whole-genome association and population-based linkage analyses,” The American journal of human genetics, vol. 81, no. 3, pp. 559–575, 2007.

[35] J. Tyrrell, S. E. Jones, R. Beaumont, C. M. Astley, R. Lovell, H. Yaghootkar, M. Tuke, K. S. Ruth, R. M. Freathy, J. N. Hirschhorn, et al., “Height, body mass index, and socioeconomic status: mendelian randomisation study in uk biobank,” bmj, vol. 352, 2016.

[36] D. A. Lawlor, “Commentary: Two-sample mendelian randomization: opportunities and challenges,” International journal of epidemiology, vol. 45, no. 3, p. 908, 2016.

[37] D. J. Weiner, E. M. Wigdor, S. Ripke, R. K. Walters, J. A. Kosmicki, J. Grove, K. E. Samocha, J. I. Goldstein, A. Okbay, J. Bybjerg-Grauholm, et al., “Polygenic transmission disequilibrium confirms that common and rare variation act additively to create risk for autism spectrum disorders,” Nature genetics, vol. 49, no. 7, pp. 978–985, 2017.

[38] N. M. Laird and C. Lange, “Family-based designs in the age of large-scale gene-association studies,” Nature Reviews Genetics, vol. 7, no. 5, pp. 385–394, 2006.

[39] G. R. Abecasis, L. R. Cardon, and W. Cookson, “A general test of association for quantitative traits in nuclear families,” The American Journal of Human Genetics, vol. 66, no. 1, pp. 279–292, 2000.

[40] N. M. Laird, S. Horvath, and X. Xu, “Implementing a unified approach to family-based tests of association,” Genetic Epidemiology: The Official Publication of the International Genetic Epidemiology Society, vol. 19, no. S1, pp. S36–S42, 2000.

[41] G. Thomson, “Mapping disease genes: family-based association studies.,” American journal of human genetics, vol. 57, no. 2, p. 487, 1995.

[42] R. S. Spielman, R. E. McGinnis, and W. J. Ewens, “Transmission test for linkage disequilibrium: the insulin gene region and insulin-dependent diabetes mellitus (iddm).,” American journal of human genetics, vol. 52, no. 3, p. 506, 1993.

[43] J. A. Labrecque and S. A. Swanson, “Interpretation and potential biases of mendelian randomization estimates with time-varying exposures,” American journal of epidemiology, vol. 188, no. 1, pp. 231–238, 2019.

[44] J. Pearl, Causality: Models, Reasoning and Inference. Cambridge University Press, 2nd ed., 2009.

